# Combi-Seq: Multiplexed transcriptome-based profiling of drug combinations using deterministic barcoding in single-cell droplets

**DOI:** 10.1101/2021.09.16.460212

**Authors:** L Mathur, B Szalai, R Utharala, M Ballinger, JJM Landry, M Ryckelynck, V Benes, J Saez-Rodriguez, CA Merten

## Abstract

Anti-cancer therapies often exhibit only short-term effects. Tumors typically develop drug resistance causing relapses that might be tackled with drug combinations. Identification of the right combination is challenging and would benefit from high-content, high-throughput combinatorial screens directly on patient biopsies. However, such screens require a large amount of material, normally not available from patients. To address these challenges, we developed a scalable microfluidic workflow to screen hundreds of drug combinations in picoliter-size droplets using transcriptome changes as a readout for drug effects. We devised a deterministic combinatorial DNA barcoding approach to encode treatment conditions, enabling the gene expression-based readout of drug effects in a highly multiplexed fashion. We applied our method to screen the effect of 420 drug combinations on the transcriptome of K562 cells using only ∼250 single cell droplets per condition, to successfully predict synergistic and antagonistic drug pairs, as well as their pathway activities.

## Introduction

Despite major progress over the last decades, cancer remains a major cause of death. Our increased molecular understanding of the molecular basis of cancer has led to the development of targeted therapies. These therapies have so far provided limited efficacy and only in a small subset of patients^1^, despite major efforts to characterize patients genomically to find response biomarkers.

An approach that holds the promise to improve this situation is to complement large-genomic profiling in basal conditions with measurements after perturbing cancer cells with drugs^2^. While many approaches can be used to perform drug screenings, they are often low in throughput^3^, cost and time extensive^4^ and/or require large amounts of cells^5^, which together strongly restricts the number of potential drugs that can be screened per tumor biopsy. This limitation gets more pronounced when considering drug combinations due to the sheer number of potential combinations, which increases exponentially with the number of tested drugs.

Due to limited screening capacities, computational approaches to model drug-drug interactions have been developed^6^. While models on drug efficacies improved over the past years by an increase in available data resources, predictions on drug responses remain challenging and limited to well characterized systems such as cell lines, thereby limiting their translatability into clinics. Among the different data types, gene expression states of cells were shown to be highly predictive for drug response^7^. Additionally, data repositories of drug-induced transcriptional changes, such as LINCS^8^, have proven to be a valuable resource. While there are already perturbation screening platforms available in plates for bulk^9,10^ and single cell^11,12^ transcriptomics, they usually require large numbers of cells per tested condition, and they have not been used for screening drug combinations. Therefore, integrating transcriptomic readouts into a miniaturized combinatorial drug screening platform with the potential to screen tumor biopsies will enable more relevant predictions and increase our understanding on the mode of action of synergistic and antagonistic drug-drug interactions.

Droplet based microfluidics, which uses picoliter to nanoliter sized droplets as reaction vessels to perform cellular screens, provides a promising approach to achieve this goal. Due to the miniaturization over several orders of magnitude as compared to conventional plate-based screens, the number of drugs or drug combinations can be massively upscaled while working with low input cell numbers^13^. We previously demonstrated a first step in this direction by integrating Braille valves into a droplet microfluidic system to generate drug combinations in so called plugs (∼500 nl large droplets) stored sequentially in tubings^14^. Plugs were used to directly screen 56 combinatorial treatment options on pancreatic tumor biopsies to find the most potent drug pairs using a phenotypic apoptosis readout. While our previous approach provided a first proof of concept in directly screening patient material, the still relatively large volumes of 500 nl limited the number of drug pairs tested. Furthermore, an apoptosis assay provides only a single endpoint readout with limited insights into the drug pairs’ mode of action, which could significantly improve our understanding and the predictability of drug combinations to tackle resistance mechanisms.

To overcome these limitations, we present here a microfluidic platform that allows to perform highly multiplexed screens of hundreds of drug combinations in an emulsion of picoliter sized droplets. By introducing a deterministic combinatorial barcoding approach, where sets of two barcodes encode drug pairs, we managed to screen all conditions in a highly multiplexed fashion, without the need to keep any spatial order (e.g. wells, plug sequence). Since the DNA barcodes were designed for whole transcriptome analysis of cells after drug perturbation, we were additionally able to perform massively parallelized gene expression-based profiling of drug combinations. We demonstrated that the presented approach can be applied to determine the impact of drugs on cell viability and cellular signaling, thus providing a high throughput approach to discover synergistic drug pairs and to decipher their mode of actions.

## Results

### Microfluidic workflow to generate drug combinations in picoliter sized droplets

Multiplexed combinatorial drug screens were performed in single cell droplets by encapsulating drugs together with DNA barcode fragments (**Fig. 1a**). Each pairwise drug combination was encoded by a unique combination of two DNA barcode fragments, which together provided a priming site for reverse transcription (poly-dT) and PCR. After off-chip incubation of droplets, reagents for cell lysis, barcode fragment ligation and reverse transcription were added to each droplet by picoinjection^15^. The ligation of two barcode fragments (BC-RT and BC-PCR) resulted in functional barcodes, encoding pairwise drug combinations (**Fig. S1**). Since the barcodes were used for the reverse transcription of mRNA released from lysed cells, transcriptomes were barcoded according to drug treatments (**Fig. 1b**). Subsequently, barcoded cDNA was extracted from the droplets to construct a sequencing library. Finally, sequencing was performed to demultiplex treatment conditions and to analyze their effects on gene expression.

**Fig. 1.**
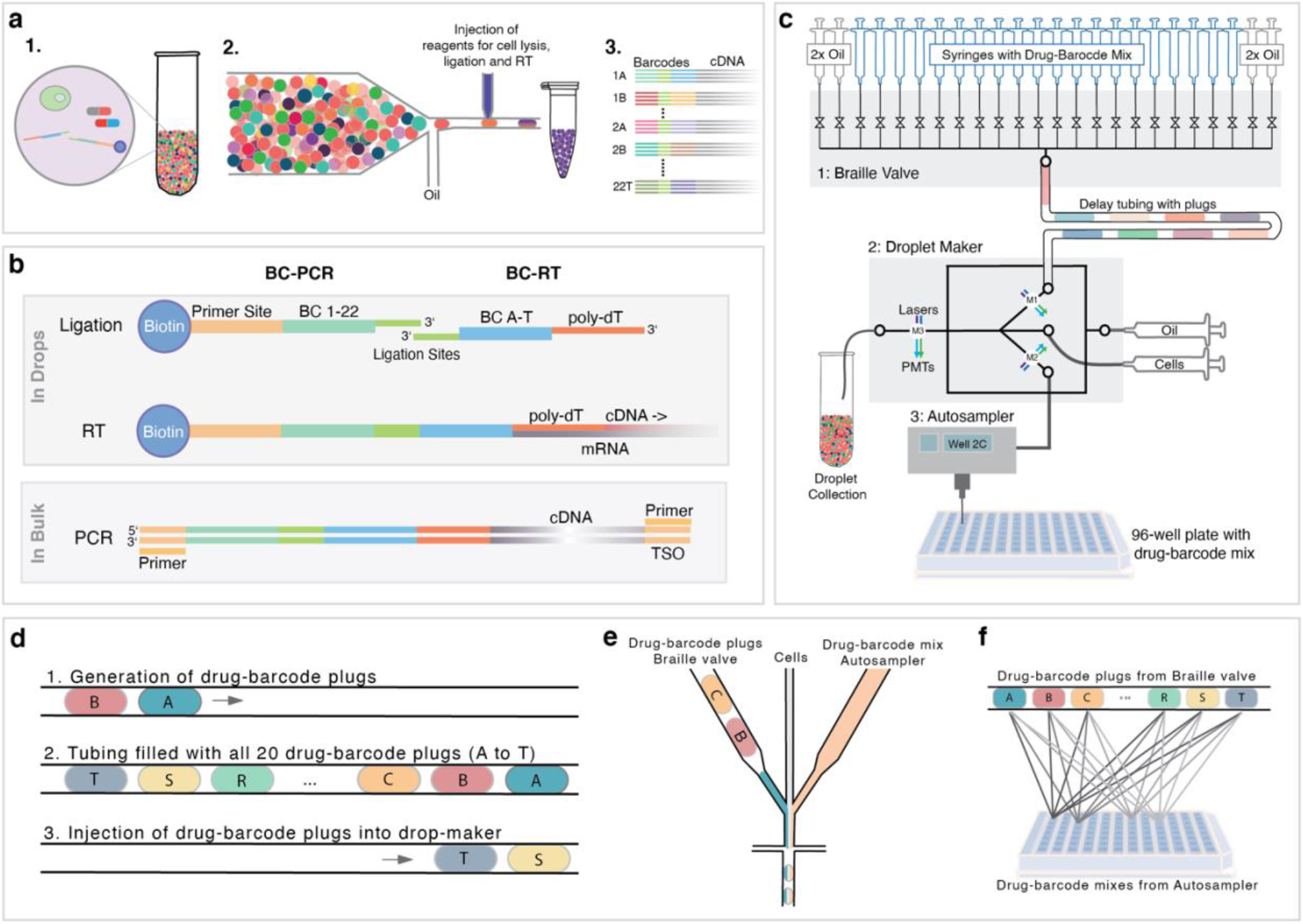
Workflow overview and microfluidic pipeline: **(a)** Workflow of the combinatorial drug screen with gene expression based read out. (1) Cells were encapsulated along with drug combinations and pairs of barcode fragments encoding drugs. (2) After 12h off-chip incubation at 37 °C, droplets were reinjected into a second chip and pico-injection was used to add reagents for cell lysis, barcode ligation and reverse transcription (RT), enabling barcoding of the transcriptome according to the drug treatment. (3) Barcoded cDNA libraries were generated for sequencing which facilitated demultiplexing of drug treatments and gene expression-based readouts. **(b)** Barcoding strategy applied to encode and decode drug combinations. Pairs of barcode primers consisting of a biotinylated barcoded PCR primer (BC-PCR) and a barcoded poly-dT primer (BC-RT) were joined in a ligation reaction to form functional barcodes encoding a combination of two drugs. Reverse transcription of the mRNA incorporates a barcode combination encoding the perturbation the cells were exposed to into each transcriptome. By breaking droplets, barcoded cDNA can be recovered over the biotin handle and amplified for sequencing using primer sites on the BC-PCR and a template switching oligonucleotide (TSO). **(c)** Scheme illustrating the microfluidic pipeline used to generate drug combinations in droplets. A Braille valve module (1) was used to direct the injected drug-BC-RT mixes and oil either to a waste outlet or into a delay tubing. By opening each valve sequentially and by injecting oil between each drug, a sequence of 20 drug-barcode plugs was generated within the delay tubing. The delay tubing was connected to a drop maker (2) into which cells were injected using a syringe pump and drugs and BC-PCR fragments from a 96-well plate were co-injected by an autosampler. Finally, injecting the plugs into the drop maker by opening two oil valves, droplets containing drug pairs with barcodes and cells were generated. Labels M1 – M3 show at which positions fluorescence signals were measured (e.g. for plots in figure 2) **(d)** Generation of drug-barcode plugs in the delay tubing: (1) Plugs spaced out by oil were produced by sequentially opening the corresponding valves, (2) resulting in a delay tubing filled with a sequence of 20 drug-barcode plugs. (3) By opening two oil valves, the plugs in the delay tubing were injected into the droplet-maker chip. **(e)** Droplet production using an aqueous phase consisting of cell suspension, drug-barcode plugs and drug barcode mixtures injected by the autosampler, resulting in the co-encapsulation of cells together with drug-barcode combinations from the valve module and the autosampler. **(f)** Scheme illustrating the generation of drug combinations from 20 drugs from the valve module with drugs from a 96-well plate. Each sequence of 20 drug-barcode (BC-RT) plugs was combined with drug-barcode (BC-PCR) mixes from one well.

In order to generate drug combinations in picoliter sized droplets, we synchronized the Braille valve system and autosampler based injection of drugs into a droplet-maker chip **(Fig. 1c**). In addition, cell-suspensions were injected into the droplet-maker chip at a density of 0.1 cells per droplet volume, to obtain droplets containing single cells (**Movie S1**). The autosampler (Dionex) was loaded with a 96-well plate, with each well containing a single drug together with the corresponding barcoded primer fragment (BC-PCR) and a marker dye enabling to monitor later mixing steps. Drugs were consecutively aspirated and injected into the droplet-maker. The time window between two samples from the autosampler (∼3 min) was used to generate a sequence of 20 chemically-distinct plugs, each containing unique pairs of two drugs and two barcode fragments (BC-RT and BC-PCR), by injecting secondary drugs and barcodes (BC-RT) into a separate Braille valve chip (**Fig. 1c** and **Fig. S2a**). In particular, each compound valve was opened sequentially and fluorinated oil was injected in between, so that drug-barcode plugs spaced out by an immiscible oil phase could be injected into a delay tubing (**Fig. 1d**). Once the delay tubing was filled with a sequence of 20 plugs, two oil valves were opened to inject all plugs into the droplet maker (∼2 min, **Fig. S2b**). Thereby, drug-barcode plugs from the valve system were combined with the drug-barcode mixtures being injected from the autosampler and encapsulated together with single cells into droplets (**Fig. 1e**). By repeating this process, hundreds of combinations with specific pairs of barcode fragments were generated (**Fig. 1f**). It is important to note that scaling up the number of combinations can be achieved by increasing the number of drug injected from the autosampler.

Synchronization between the autosampler and the valve-based injection of drugs was crucial to ensure that combinations were only generated once the drug injected from the autosampler had reached its plateau concentration. Between each drug coming from the autosampler, a time window with decreasing and increasing concentrations was observed, as shown by the alternating injection of fluorescence dyes (**Fig. 2a**). This phenomenon is based on Tailor-Aris dispersion of solutes in the continuous, miscible carrier phase (PBS) of the autosampler^16^.No combinations were generated during that time window, which was rather used to produce compound plugs in the delay tubing of the Braille module. Once the plateau concentration was reached, as indicated by measuring a constant intensity of a fluorescent marker dye, the 20 compound plugs were injected into the droplet-maker and combined with the drug from the autosampler and cells into droplets. The injection of one such plug train took 2 min, resulting in an overall time (plug production and injection) of 15 seconds for generating ∼2500 droplets containing cells and one barcoded combinatorial treatment condition. Once all 20 plugs were injected, the autosampler started aspirating the subsequent drug.

**Fig. 2:**
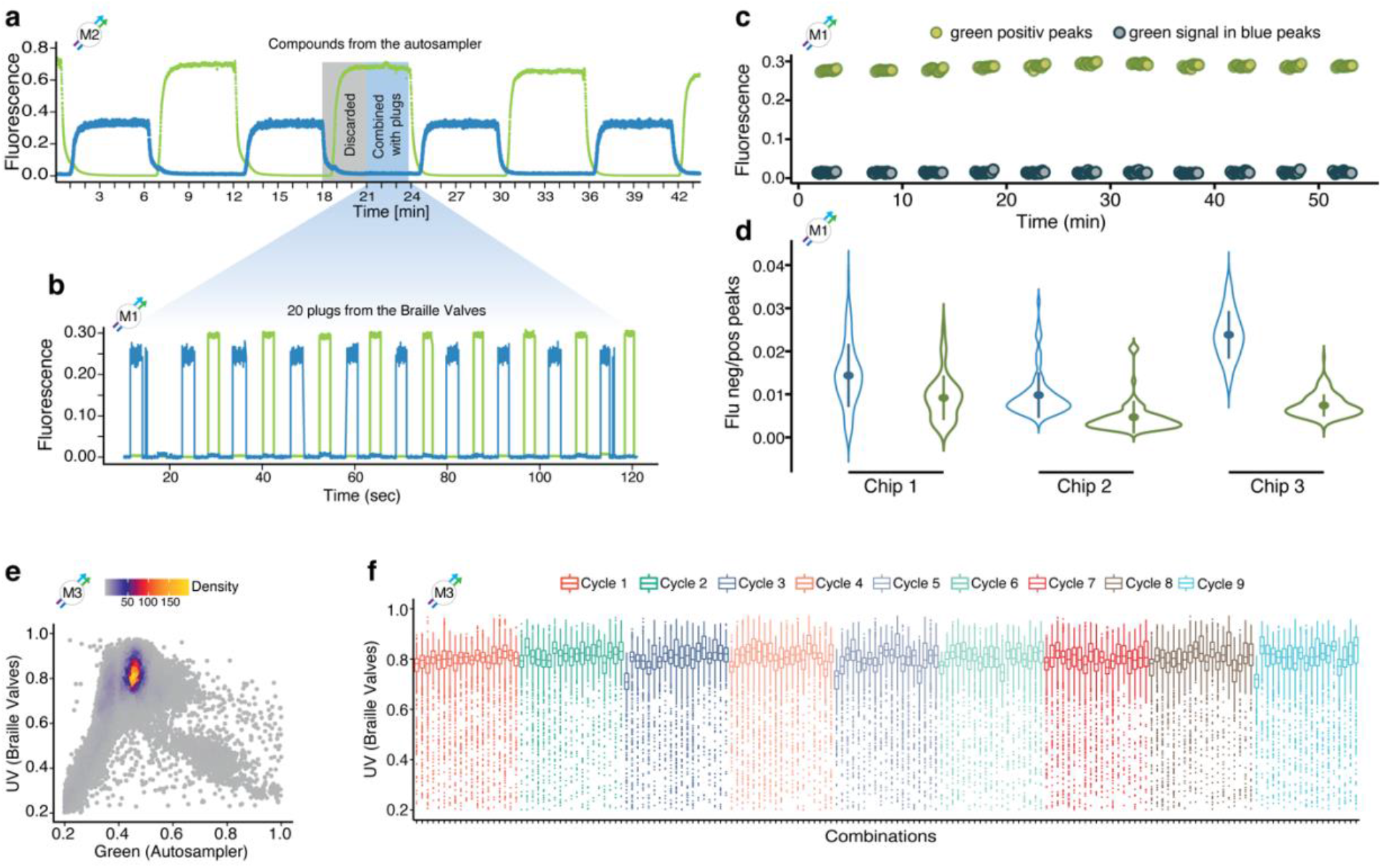
Validations of the microfluidic pipeline: **(a)** A sequence of compounds supplemented alternatingly with Alexa-488 or Cascade blue was injected from a 96-well plate using the autosampler. Samples being injected in the time window of decreasing and increasing fluorescence signals (grey box) were discarded by ensuring no combinations were generated, whereas time windows with stable fluorescence signals (blue box) were used to generate drug combinations by the co-injection of 20 plugs generated on the Braille display chip. Since no drug-barcode plugs from the Braille display were injected during time windows with instable concentrations (grey box), droplets only contained a single barcode (BC-PCR) and so that mRNA from droplets produced during these time windows was not reverse transcribed and hence discarded. (**b)** Fluorescence intensities from a sequence of plugs from the Braille valves supplemented with either Alexa-488 or Cascade Blue measured at the delay tubing inlet of the droplet marker (= before combinatorial mixing). The blue overlay connecting (a) and (b) illustrates a time series during which a cycle of 20 Braille display plugs (2^nd^ plug without reference dye) is combined with one sample from the autosampler. The alternating sequences of drugs supplemented with Cascade Blue or Alexa-488 were used to quantify cross-contamination between specific (e.g. green) fluorescent-positive plugs into the subsequent negative (e.g. blue) sample for both injection modes, separately. **(c)** Measured cross-contamination from green positive plugs into green negative plugs over a total of 209 plugs injected from the Braille display (11 cycles as shown in b). **(d)** Cross contaminations in the plugs coming from the Braille display for three different chips: The ratio between fluorescence intensities of a blue or green negative plug and the previous blue or green positive plug was analyzed to quantify the level of cross-contamination between sequential samples. **(e)** Fluorescence signals of droplets after generating combinatorial mixtures. Scatterplot representing the fluorescence intensities measured for droplets generated from only Cascade Blue labelled compound plugs from the Braille valves and only Alexa-488 labelled compound injected from the autosampler. Fluorescence signals were measured at the droplet outlet. One highly dense population of double positive droplets was observed. **(f)** UV fluorescence intensities measured for individual droplets from 180 combinations. Each color represents one cycle of 20 drug plugs combined with one drug from the autosampler. Fluorescence intensities were measured at positions M1 – M3, as indicated on the top left of each figure and in Fig. 1c. Plots e) and f) show a total of 91899 droplets, each.

To ensure droplet contents with marginal cross-contamination, we designed the geometry and delay tubing connectors of the Braille valve drop-maker chips such that no residual drug-barcode mixtures remained in the channels (**Fig. S3**). Before each experiment, we measured a proxy for the level of contamination between plugs from the Braille display. This was done by using drugs supplemented alternatingly with Alexa-488 or Cascade Blue, resulting in an alternating sequence of blue and green fluorescence peaks (**Fig. 2b**). Fluorescence intensities of plugs were measured on the droplet maker and contaminations of drugs/dyes from one plug into the subsequent plug were detected by either green signals in UV peaks or UV signals in green peaks (**Fig. 2c**). The ratio between the fluorescence signals from each negative peak (n) in either the green or the UV channel with the previous positive peak (n-1) was used as a proxy to quantify the level of cross contaminations between two drugs (**Fig. 2d**). Over three different chip setups (Braille valve and drop-maker), we found a mean of 1.5% of contamination in the UV channel and 0.7% in the green channel (**Table S1**), indicating that the described systems can be applied to generate combinations with sufficiently high purity.

In the described microfluidic pipeline, drug combinations were generated by mixing drugs injected from an autosampler and a Braille valve module. To ensure the precise and accurate drug concentrations within droplets, both drugs had to be encapsulated at a constant and predefined ratio. We validated the precise mixing of two drugs by supplementing all compounds on the braille display with Cascade Blue and all compounds from the autosampler with Alexa 488. We observed one highly dense main population of double blue and green double positive droplets, demonstrating that both compounds were co-encapsulated at a constant ratio (**Fig. 2e**). Furthermore, we confirmed the stable co-encapsulation of the two dyes for droplets over individual combinations (**Fig. 2f**). The median fluorescence intensities of individual combinations were highly stable with coefficients of variation (CV) over 180 combinations of 2.9% and 3% for blue and green intensities, respectively. The scattering of droplets around the main population can be explained by a short (< 100 ms) flow equilibration phase at the beginning and end of plugs and fluctuation of the droplet trajectory within a focused laser beam (**Fig. S4, Movie S2**). Consequently, we concluded that the injections modes (Braille valve or autosampler) can be robustly synchronized to generate drug combinations in droplets at high precision and purity.

### Validations of gene expression based combinatorial drug screens

To characterize the microfluidic pipeline and to demonstrate its applicability to perform gene expression-based combinatorial drug screens, we designed a small 4×4 drug screen (**Table 1**). First, we wanted to assess whether the injection mode of drugs from the Braille valves vs. autosampler causes any bias, and therefore loaded the same set of drugs on the Braille valves and autosampler. In case of an injection bias, we would expect to see differences between the same combination generated in reverse order (e.g. Imatinib and Trametinib vs Trametinib and Imatinib). Secondly, we aimed at assessing the impact of the barcoding mode on the gene expression readout. For this purpose, we first encoded treatment conditions such that drugs injected from the Braille valves were supplemented with barcoded BC-RT, whereas drugs from the autosampler were supplemented with BC-PCR. Then we repeated the experiment with BC-RT encoding drugs from autosampler and BC-PCR encoding drugs from the braille valves, expecting comparable results if the barcoding mode is not impacting the readouts. We used the described pipeline to generate droplets each containing single human leukemia K562 cells and all pairwise combinations of drugs and the corresponding barcodes and incubated the emulsion for 12h. After ligation, the two barcoded primer fragments formed one functional barcode encoding the pairwise drug combination. In order to obtain three replicates, the whole process was performed three times.

**Table 1:** Matrix of drugs used in the combinatorial 4×4 screen

|  | Drugs | Drug Braille Valves |  |  |  |
| --- | --- | --- | --- | --- | --- |
|  |  | Imatinib | Trametinib | YM155 | DMSO |
| Drugs Autosampler | Imatinib | Imatinib<br>Imatinib | Trametinib<br>Imatinib | Imatinib<br>YM155 | DMSO<br>Imatinib |
|  | Trametinib | Imatinib<br>Trametinib | Trametinib<br>Trametinib | YM155<br>Trametinib | DMSO<br>Trametinib |
|  | YM155 | Imatinib<br>YM155 | Trametinib<br>YM155 | YM155<br>YM155 | DMSO<br>YM155 |
|  | DMSO | Imatinib<br>DMSO | Trametinib<br>DMSO | YM155<br>DMSO | DMSO<br>DMSO |

Each ligated barcode was used to reverse transcribe the transcriptomes from perturbed cells (**Fig. 1a**). After data preprocessing (Methods) and initial quality controls (Median read and gene count per sample of 3.47×10^5^ and 3229, respectively, **Fig. S5**) we performed dimension reduction using t-distributed stochastic neighbor embedding (t-SNE, **Fig. 3a**) on the demultiplexed count matrix. To analyze whether some systematic bias arises based on the injection source (autosampler or Braille valves), we analyzed samples according to the autosampler drug (**Fig. 3a**, top left panel), the Braille valves drug (**Fig. 3a**, top right panel), the “ordered” drug combination (where we made distinction between e.g.: Imatinib-Trametinib and Trametinib-Imatinib combination), and the “unordered” drug combination (where Imatinib-Trametinib and Trametinib-Imatinib samples were not distinguished). While the injection mode for single drugs from the autosampler (**Fig. 3a**, top left panel) or the Braille valves (**Fig. 3a**, top right panel) has only moderate impact on the clustering of individual data points, their pairwise combinations (**Fig. 3a**, bottom panels) is the stronger determinant on the cohesion and separation between samples. To further quantify the extent of sample clustering based on injection source for ordered and unordered combinations we performed silhouette analysis (**Fig. 3b**, and Methods for further details). As the distribution of silhouette scores are dependent on the number of clusters (4 for drugs, 16 for unordered and 10 for ordered combinations), we compared the silhouette scores of clustering to random distributions created by permuting the sample labels. The silhouette scores for single drugs and combinations were significantly higher (p values < 0.01) than the background distributions, showing that samples cluster together based on the used drugs and combinations. Consequently, also the barcoding mode for single drugs injected from the Braille valves or the autosampler encoded either with barcoded RT or PCR primers do not introduce a bias, since their impact on clustering and hence gene expression, is indistinguishable. Contrary to this, pairwise combinations were driving clustering of the samples, showing that both drugs were detected together in an unbiased way.

**Fig. 3:**
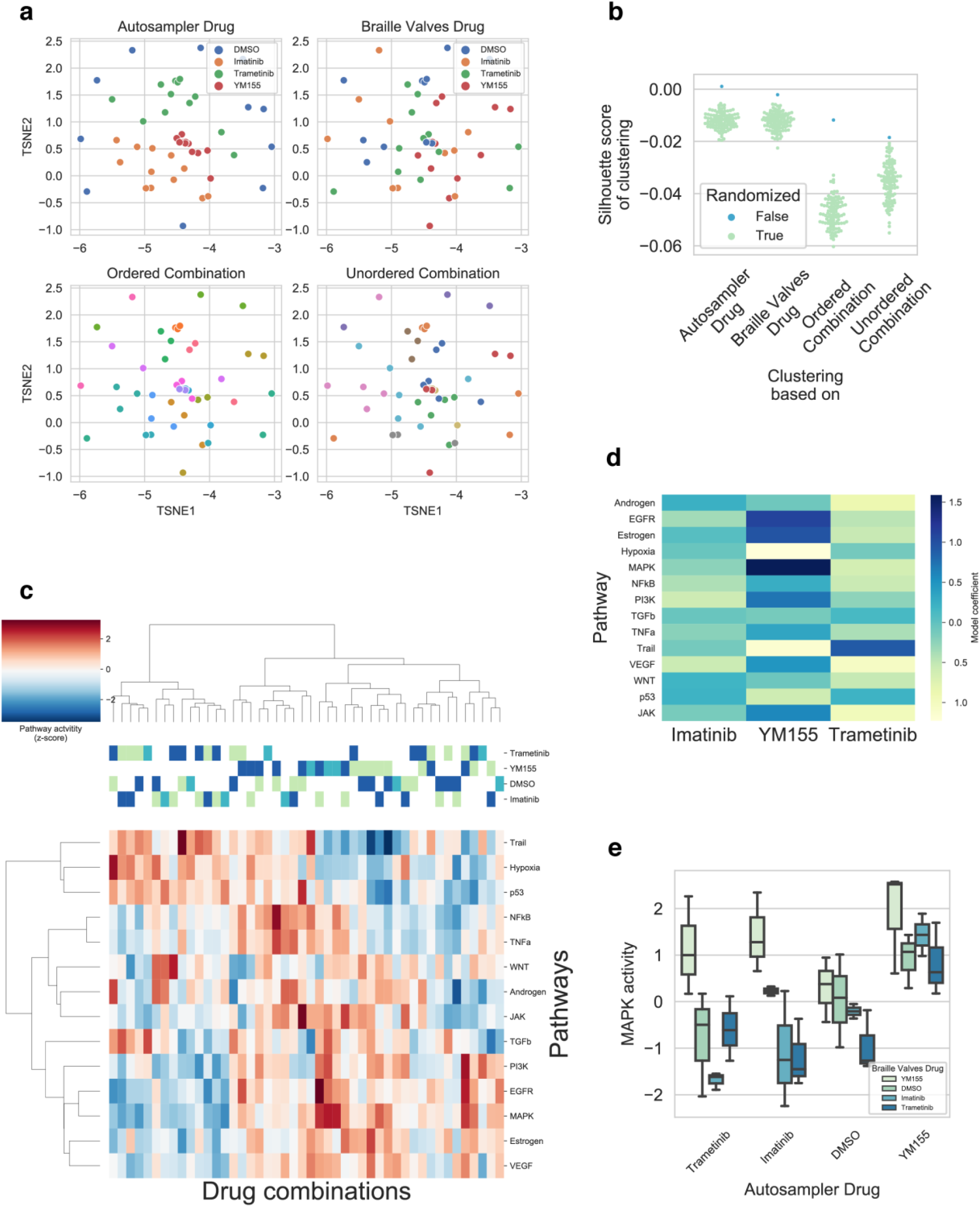
Validations of gene expression based combinatorial drug screens. **(a)** TSNE plots of normalized gene expression data. Samples are color coded based on Autosampler drug (top left panel), Braille valves drug (top right panel), ordered combination (bottom left panel) and unordered combination (bottom right panel). Color code is labeled for autosampler and Braille valves drugs (top panels). **(b)** Silhouette scores of sample clustering based on autosampler / Braille valves drugs and ordered / unordered combinations. Silhouettes scores are compared to random distributions (color code) created by permuting sample labels. **(c)** Pathway activity heatmap of samples. PROGENy pathway activities were calculated for each sample (z-scores of pathway activities, color code) and the pathway activity matrix was hierarchically clustered. Drugs of combinations are color coded (yellow: autosampler drug, blue: braille valves drug, cyan both drugs). **(d)** Drug induced pathway activity changes. A linear model (pathway_activity ∼YM155 + Imatinib + Trametinib) was fitted for each pathway, and the linear model coefficients (color code) for each drug is plotted as a heatmap. **(e)** Drug induced MAPK activity changes. MAPK activity (y axis) grouped based on autosampler drug (x axis) and Braille valves drug (color code) and plotted as a boxplot (median, quartiles and full distribution).

We also performed hierarchical clustering using the 100 most highly expressed genes across samples, which also showed drug and combination-based clustering of the samples (Fig. S6). To further demonstrate that our experimental pipeline does not introduce significant technical biases, we performed the small 4×4 screen with swapped barcodes (Braille valves drugs supplemented with BC-PCR and autosampler drugs supplemented with BC-RT). We observed similar quality (**Fig. S7**) and clustering (**Fig. S8**) of samples in this case.

To further analyze the gene expression signatures of cells treated with different combinations, we calculated pathway activity changes for each sample, using the PROGENy method^17-19^. PROGENy calculates pathway activities from gene expression data for 14 cancer related pathways. Hierarchical clustering of samples based on pathway activities (**Fig. 3c**) also showed the drug and combination-based clustering. We observed two main clusters, one corresponding to combinations including YM155, while the other was dominated by Trametinib treated samples. Analyzing the associations between pathways activity changes and drugs (**Fig. 3d**), we found a decreased activity of Hypoxia pathway in all YM155 treated samples, while all Trametinib (MAPK inhibitor) treated samples showed strong inactivation of MAPK (**Fig. 3e**, p value from linear model: 0.03), and related EGFR pathways. This pathway analysis suggests that the observed gene expression changes correspond to the known mechanism of action of the used drugs. This supports the use of our screening method to analyze combination induced gene expression changes in a high-throughput manner, enabling the characterization of drug responses in much greater detail as compared to phenotypic assays used previously^14^.

### High throughput gene expression based combinatorial drug screen

Based on the promising results of the 4×4 drug combination experiment, we performed a high-throughput screen, using a total of 420 different combinatorial treatment conditions. In order to estimate optimal drug concentrations for large scale combinatorial screens, we generated dose response curves using K562 cells to determine their GR35 values for each single drug (**Table S4**)^20^. Drugs were assigned to the Braille valves or autosampler to achieve a balanced distribution of drugs with high and low GR35 values. Drugs loaded on the Braille valves or autosampler were supplemented with BC-RT or BC-PCR, respectively. We aimed at generating 250 droplets containing a single cell for each of the 420 treatment conditions and incubated droplets for 12h at 37 °C before performing picoinjection for cell lysis, barcode ligations and RT (**Fig. 1a**). This process was performed three times to obtain replicates.

After initial preprocessing and quality control (median reads and genes per sample of 32892 and 547, respectively, **Fig. S9**), we performed the same dimensionality reduction (**Fig. 4a**) and silhouette analysis (**Fig. 4b**), as for the 4×4 screen. Again, samples clustered significantly better based on the used combination, than randomly expected (p values of silhouette scores vs. random distribution: <0.01, 0.47 and <0.01 for autosampler drug, Braille valves drug and combination, respectively.).

**Fig. 4.**
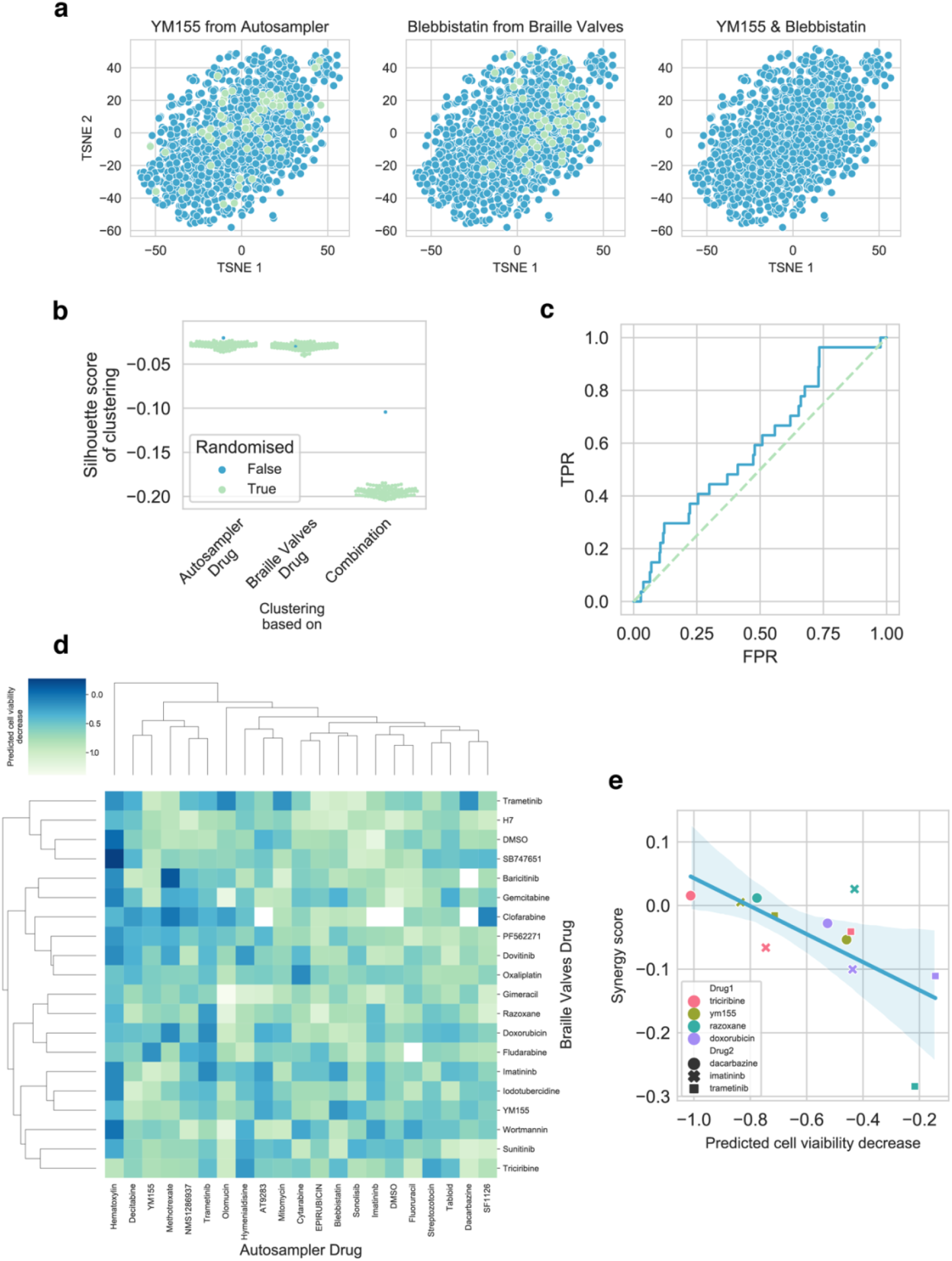
High throughput gene expression based combinatorial drug screen. **(a)** TSNE plots of normalized gene expression data. YM155 (left panel), Blebbistatin (middle panel) and YM155-Blebbistatin combination treated samples are labeled as a representative example. **(b)** Silhouette scores of sample clustering based on autosampler / Braille valves drugs and combinations. Silhouettes scores are compared to random distributions (color code) created by permuting sample labels. **(c)** ROC analysis of drug signature similarity to LINCS-L1000 data. For each drug, a consensus signature was calculated and similarity (Spearman’s rank correlation) to corresponding LINCS-L1000 signatures was calculated. The similarity values were used as predicted values for ROC analysis, while true positives were the matched drug pairs between high-throughput screen and LINCS-L1000. **(d)** Heatmap of predicted cell viability for drug combinations. Cell viability was predicted from gene expression data using the CEVIChE method. **(e)** Experimental validation of predicted cell viability. Drug synergy (y-axis) was measured for 12 combinations in a microtiter plate format (color code) and plotted against the predicted synergy value (x-axis).

To further investigate whether the gene expression values of the high-throughput screen are biologically meaningful, we compared the obtained gene expression signatures to those available for the same drugs in the public LINCS-L1000 dataset^8^. As LINCS-L1000 contains only expression signatures of monotherapy drug treatments, we calculated consensus signatures for each drug of our high-throughput screen (Methods) and compared these to consensus signatures generated from the LINCS-L1000 database across all available cell lines and concentration doses (note that LINCS-L1000 does not include data obtained directly from K562 cells). For 32 drugs used in our microfluidic screen, corresponding data on LINCS-L1000 was available. To compare signature similarities of these, we calculated Spearman’s correlation coefficients for all pairs of drugs across the two datasets. Our ROC analysis showed that signatures of the same drugs from the two screens (true positives) are more similar, than signatures of unrelated drug pairs (**Fig. 4c**, ROC AUC: 0.59), and this area under the ROC curve is statistically significant compared to a random distribution created by permuting drug labels (p=0.019).

As all used drug concentrations were GR35 values, we expected that synergistic combinations could lead to decreased cell viability, while in case of antagonistic combinations we expected increased cell viability values. While we did not measure cell viability directly, the CEVIChE algorithm (Methods)^18^ allowed us to infer cell viability changes for all used drug combinations from gene expression data (**Fig. 5c**). We found several clusters of potential synergistic and antagonistic combinations (e.g.: Triciribine-Dacarbazine and Razoxane-Trametinib, respectively). To experimentally validate these results, we performed 5×5 dose matrix combinatorial cell viability screens with all possible combinations of Triciribine, YM155, Razoxane and Doxorubicin with Dacarbazine, Imatinib and Trametinib drugs in a microtiter plate format. We calculated synergy scores (positive: synergistic, negative: antagonistic) for the tested combinations using the Bliss independence synergy model (Methods)^21^. Our measured (plate experiment) and predicted (from gene expression data obtained in the microfluidic system) synergies showed significant correlation (Pearson correlation r = -0.66, p= 0.018), further confirming the discovery potential of our platform.

In summary, using our combinatorial microfluidic gene-expression platform, we showed that i) the measured gene expression values cluster based on the chemical perturbation, ii) the resulting data is in good agreement with public monotherapy perturbation profiles and iii) predicted cell viabilities and drug synergies could be validated in a microtiter plate format for selected hits. Taken together, this illustrates how comprehensive information can be gained from gene expression profiles obtained in a highly multiplexed microfluidic format, sequencing only about 250 cells per drug treatment. This should make the workflow particularly interesting for use with very limited material, such as patient samples.

## Discussion

Cancer patient stratification for personalizing treatments with chemotherapeutics and targeted drugs have shown to increase the successes of cancer therapies^22-24^. These efforts are largely driven by dissecting the genomic and transcriptional landscape of tumors or cell lines in order to identify traits that explain drug sensitivities^25,26^. While a variety of identified genomic and/or transcriptomic markers are successfully used in clinics, they are available for only a small subset of tumor types and patients^1^. Furthermore, many patients often suffer from tumor relapse^27^, which is largely rooted in intra-tumor heterogeneity^28^. The relapse is often driven by the surge of a resistance mechanism to the drug that renders the efficacy of single drugs short lived^29^. While treatments with drug combinations offer the potential to reduce the risk of drug resistance, their prediction and empirical evaluation remains challenging.

To advance in solving these challenges, we present here a microfluidic pipeline enabling highly multiplexed combinatorial drug screens in single-cell droplets using global transcriptomics as a readout. By integrating deterministic barcoding of treatment conditions, we were able to assess the efficacy of drug combinations by changes in gene expression and gained comprehensive readouts from whole transcriptome sequencing. We applied our pipeline to screen 420 combinatorial treatment conditions in a single tube, illustrating the high level of multiplexing. Based on assay miniaturization in a droplet format, only about 250 cells were needed per tested condition, hence opening a way for personalized screens on patient material.

We have designed the microfluidic platform as a modular system in which the Braille display valves allow us to quickly change between injected drugs overcoming the limitations from a slow autosampler based injection. Since both are combined on the droplet generator, fast and efficient generation of drug combinations becomes feasible and allows the encapsulation of single cells into droplets of high chemical diversity. Since the autosampler used here injects drugs from up to three 96 or even 384-well plates, the number of drug combinations can be further scaled up to a theoretical maximum of 3 × 384 × 20 = 23040 combinations in a single experiment. What becomes most limiting at that scale are sequencing costs and available material (when e.g. using primary cells) rather than instrument throughput.

We see significant added potential by the possibility to screen such large numbers of drug combinations at the single cell level: Integrating fluorescence-based droplet sorting upstream of the picoinjection (cell lysis) step could, for example, be used to physically separate and sequence resistant clones for all 420 treatment options in a single experiment (e.g. implementing the phenotypic Caspase-3 assay we used previously)^14^. This way one could analyze the difference in their transcriptomic signature as compared to responding cells, opening the way for highly multiplexed studies to reveal new biomarkers for resistance and (chemical) sensitizers to overcome these. As recently demonstrated, single cell readouts of drug perturbation provide great insights into heterogeneous drug response^11^. Performing such screens on patient biopsies will allow us to dissect subclonal drug responses and thereby to define more efficacious drug combinations and additionally gain insights into their potential resistance mechanisms. In order to enable the encapsulation of primary tumor cells, we aim at integrating support structures for cell adherence and growth in droplets as previously shown^30,31^.

All generated datasets demonstrate that neither the barcoding approach nor the injection mode biased the gene expression-based readout. Monotherapies from both injection and barcoding modes had similar impacts, while their combinations had the strongest impact on gene expression and was the main driver of the observed clustering. This confirms the highly precise and accurate operation of the presented microfluidic workflow and the specificity of the deterministic barcoding approach. Additionally, we found significant similarities between consensus gene expression signatures of monotherapies from our large-scale screen with drug signatures from the LINCS-L1000 dataset, illustrating a high level of reproducibility. In the pathway activity analysis, we found that the hierarchical clustering was largely driven by the three drugs YM155, Imatinib and Trametinib, which further supports the detection of drug-specific effects. The two main clusters were driven by Trametinib treatments inhibiting MAPK and EGFR pathway activities, and the opposing effects of YM155 treatments, inducing the up-regulations in MAPK and EGFR pathway activities while inhibiting hypoxia and TRAIL related pathways. While the effects of Trametinib on MAPK are expected^32^, the effects of survivin inhibition by YM155 are less well understood, due to complex signaling and incomplete knowledge on survinin^33^. The increased MAPK pathway activity is likely to reflect a counteractive mechanism, since survivin expression was described to be regulated by Sp1 and c-Myc activation through the MAPK pathway^34^ and higher concentration of the drug target will reduce the drug effect. As survinin expression has been linked to drug resistance in leukemia, a combinatorial treatment with YM155 and Trametinib could potentially have a beneficial effect on decreasing the chances for relapse, due to the inhibition of survivin and putative compensatory expression induced by the MAPK pathway. Taken together, these findings show that the described microfluidic pipeline can be applied to disentangle the effects of drug combinations on pathway activities. Such information will be of great impact when screening patient biopsies to identify potential resistance mechanisms and to predict efficacious drug pairs. Analyzing the pathway activities upon perturbations was limited by the number of detected genes, and therefore, to the small 4×4 screens, since these samples were sequenced at higher depth. To detect a comparable number of genes for all 420 drug combinations, a ten times higher coverage would have been necessary. We instead used the large screen to show that shallow sequencing data is sufficient to determine synergistic drug pairs. We mined the dataset with 420 treatment conditions for drug pairs with synergistic or antagonistic effects and identified Triciribine-Dacarbazine and Razoxane-Trametinib combinations as corresponding examples. Furthermore, we validated that there is good correlation between viability scores from the gene expression data and the experimentally validated synergy scores obtained from the plate-based drug screen. These results show that it is possible to use cost effective low sequencing depth in large transcriptomics screens to discover synergistic drug pairs.

Together with the inferred pathway activities under perturbation, this should not only allow for the identification of synergistic combinations, but also gain insights into their mechanisms of action. Compared to our previous single-measurement phenotypic assay platform^14^, the global transcriptomic readout provides orders of magnitude more data points per sample, while the cell consumption could be reduced further by a factor of about 6-fold. The higher content readouts should enable more robust predictions on the best combinatorial treatments and the discovery of new drug sensitizers and biomarkers, and the even smaller needs of material further facilitate the application in the clinic for patient stratifications and treatment prioritization.

## Methods

### Braille valve module

All devices used for the valve module were replicated from moulds prepared using soft-lithography with AZ-40XT positive photoresist (Microchemicals) according to the manufacturer’s instructions. Structures from 25400 dpi photomasks (Selba) were patterned on 4-inch silicon wafers (Siltronix) in a mask aligner (Suess MicroTec MJB3) using light with a wavelength of 375 nm. Structures were covered with a ∼1 cm thick layer of PDMS mixed with curing agent at a 1:10 ratio (Sylgard 186 elastomer kit, Dow Corning Inc) and cured overnight at 65 °C. In addition, we prepared PDMS membranes by mixing PDMS with curing agent at a 1:10 ratio and distributing it over a transparent sheet using a spin coater at 500 rpm (Laurell WS 650), which were cured overnight at 65 °C. The drug inlet and waste ports of the valve chip were punched using 0.75 mm biopsy punches (Harris Unicore), whereas the plug outlet port was punched horizontally to the outlet channels using a 0.5 mm biopsy punch (Harris Unicore). Chips were bonded to a PDMS membrane using a plasma oven (Diener Femto). We inserted PTFE tubings with an inner diameter of 0.4 mm (Adtech) into the horizontally punched outlet port until the tubing reached the funnel-like structure of the outlet channel. Subsequently, chips were bound to a glass slide to support chip structures with inlets and outlets. In order to prevent surface wetting, channels were treated with Aquapel (PGG Industries) before use. The valve structures of Braille chips were aligned (Fig. S2a) on top of the pins of a Braille display (KGS Corporation, Fig. S10a) and mounted using a plexiglas holder (Fig. S10b). Using our “SamplesOnDemand” LabVIEW software (all required software can be downloaded from www.epfl.ch/labs/lbmm/downloads/mathur). the movement of the pins was controlled, so that opening the collection channel results in closing of the waste channel and vise versa. Closing a channel was achieved by a pin pushing into the elastic PDMS membrane. This could be opened again by moving the pin down. For all experiments, we used 20 syringes (Becton Dickinson) filled with 5 ml drug-barcode solutions and 4 syringes filled with 5 ml HFE oil (Novec™ 7500, 3M). These were connected to the inlet ports of the valve chip with PTFE tubing, and fluids were injected at 500 µl h^-1^ using syringe pumps (Harvard Apparatus). The waste outlets were connected with a piece of PTFE tubing to direct fluids to a waste container.

### Preparing a drop maker

Drop maker moulds were manufactured from negative photoresist SU-8 2075 as described by the manufacturer (Microchemicals). PDMS containing 10 % (w/w) curing agent was poured over the moulds and cured overnight. Inlet ports for cells and HFE were punched vertically using 0.75 mm or 1 mm biopsy punches for PEEK tubing coming from connecting the autosampler. Inlet ports for compound plugs from the Braille valves were punched horizontally using 0.5 mm punches. Chips were first plasma bound to a PDMS membrane and subsequently to a glass slide. The channel wall hydrophobicity was increased by injecting Aquapel (PGG Industries) into the channels.

### Autosampler operations

To facilitate the injection of drugs from microtiter plates into the drop maker device, we used a Dionex 3000SL Autosampler aspirating drugs from a 96-well plate. The autosampler was programmed to sequentially aspirate 310 µl of compound from wells into a 125 µl sample loop. The large excess of aspirated volume accounted for a needle volume of 60 µl and a loop overfill factor of 2. By overfilling the sample loop by twice its volume, we ensured that the remaining compound mixture was not diluted by the carrier fluids that remain in the sample loop after each cycle due to washing. The injection of compounds from the sample loop into the droplet maker was driven by a syringe pump (Harvard Apparatus) injecting PBS (Thermo Fisher) at 500 µl h^-1^. After the aspiration of one drug, a delay time started to ensure that each drug got injected and combined with all drugs from the Braille display, before the next drug was aspirated.

### Deterministic combinatorial barcoding system

Random 10 nt long DNA-sequences with balanced base distributions were generated using the bgen tool (gear.embl.de). Barcoded PCR primers were functionalized with a 5’-end biotin for purification, followed by a spacer sequence, a common primer sequence, a unique barcode sequence and a ligation site (Tab. 2). The reverse complements (RC) were functionalized with a free 5’-end phosphate to enable ligation. Barcoded RT primers comprised a dT(20)-VN sequence, a unique barcode sequence and a phosphate group at the 5’-end. The RC for this had a ligation site complementary to the ligation site of the PCR primers. A list of all barcode sequences can be found in the supplementary materials. Complementary sequences were annealed at equimolar concentrations by heating mixtures to 95 °C for 10 min in a thermal block (Eppendorf) followed by their cooling to room temperature (RT) for 1h.

**Tab. 2:**
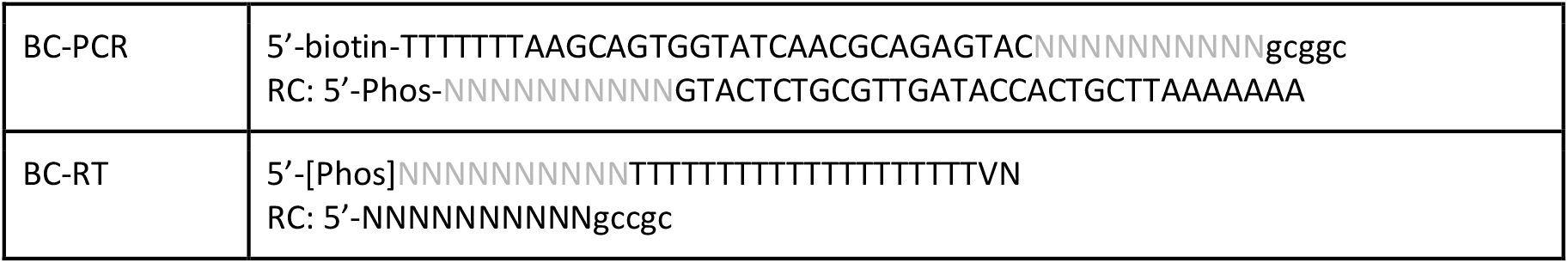
Sequences used in the barcode fragments

### Barcode-drug mixtures used for the Braille valves and autosampler

Barcode-drug mixtures for the valve module were prepared by diluting barcoded RT primers in FreeStyle media (ThermoFisher) to 1 µM. Drugs dissolved in DMSO were added to their corresponding barcode at 2x the final concentrations (see supplementary materials). The barcode-drug mixtures were supplemented with either Cascade Blue (ThermoFisher) or Alexa-488 (ThermoFisher) at 10 µM for monitoring purposes and subsequently aspirated into 5 ml luer-lock syringes (BD) connected with PTFE tubings using 27G ¾ needles (BD). Barcode-drug mixtures for the autosampler-based injection were prepared in round bottom 96-well plates by diluting barcoded PCR primers to 4 µM in FreeStyle media and the corresponding drugs to 4x the final concentrations. Mixtures were supplemented with Alexa-488 at 10 µM and plates were sealed with adhesive qPCR seals (ThermoFisher).

### Preparation of cell suspensions

K562 cells (ATCC) were cultured in IMDM media (ThermoFisher) supplemented with 10% FBS (ThermoFisher) and 1% Penicillin-Streptomycin (ThermoFisher). On the day of the experiments, cells were washed twice in PBS and resuspended in FreeStyle Media (ThermoFisher) supplemented with 4% FBS. The concentration of the cell suspension was adjusted to 2×10^6^ cells ml^-1^ and subsequently aspirated into a 3 ml luer-lock syringe (BD).

### Operations of the microfluidic pipeline for combinatorial drug screens

Syringes containing drug-barcode mixtures and HFE oil were connected to the Braille valve chip as shown in figure 1C and all injected at 500 µl h^-1^. The default mode for all compound valves was to direct fluids to the waste outlets and two HFE oil valves to direct fluids to the outlet tubing. The length of outlet tubing of the Braille valve was adjusted to harbor all 20 compound plugs spaced out with HFE oil and then connected to the Braille inlet on the drop marker chip (Fig. S2b). A syringe containing cells was mounted on a pump and connected to the drop maker and injected at 500 µl h^-1^. Cell sedimentation was prevented by low-speed rotations of the magnetic disc using the Multi Stirrus™ system (VP Scientific). Autosampler output tubing and HFE carrier phase supplemented with 1% Pico-Surf1 (Sphere Fluidics) were connected via the respective inlets and injected at 500 µl h^-1^ and 6000 µl h^-1^, respectively. The droplet maker chip was mounted on a microscope (Nikon) with an optical setup for measuring fluorescence intensities of compound plugs or droplets as described previously^14^. Briefly, lasers with a wavelength of 375 nm or 488 nm were used to excite dyes and emitted light (450 nm or 520 nm) was measured using photomultiplier tubes.

A CSV file containing the Braille valve opening sequence (**Tab. 3**) was loaded into the sample on demand software. The number of cycles was set to 21, since we were combining all 20 drugs from the Braille valves with 21 drugs from a 96-well plate. Once all tubings were connected and fluids were injected into the drop marker (carrier fluid from the autosampler and HFE oil from the Braille valves), plug production and autosampler-based injection were started simultaneously. 20 plugs were produced into the delay tubing (180 sec) and subsequently injected into the drop-maker, by opening two HFE oil valves (tot. flow rate of 1000 µl h^-1^). This ensured a continuous and stable flow rate for plug injections, and therefore, resulted in a laminar flow of compounds from the Braille valves, autosampler compounds and cells from which droplets were generated at the flow focusing junction (Movie S1). Droplets of approx. 800 pl were collected in an Eppendorf tube which was kept on ice. Once 420 combinations were generated (∼100 min) the Eppendorf tube was placed in a humidified incubator at 37 °C and 5% CO_2_ atmosphere to incubate cells for 12h.

**Tab. 3:** Braille valve operations

| Valves to open | Opening times | Outcome | Number of cycles |
| --- | --- | --- | --- |
| A | 7 sec | Compound Plug | 21x<br>Can be adapted to the number of drugs in the 96-well plate |
| 2x oil | 2 sec | Oil Spacer |  |
| B | 7 sec | Compound Plug |  |
| 2x oil | 2 sec | Oil Spacer |  |
| 17x |  |  |  |
| T | 7 sec | Compound Plug |  |
| 2x oil | 120 sec | Plug injection |  |

### Picoinjection for cell lysis, barcode ligation and reverse transcription

Chips for picoinjection were produced by replicating SU-8 moulds using PDMS with curing agent as described above. Casts were plasma bound to glass slides and treated with Aquapel. Chips were heated to 95 °C and first low melting solder and second cables were inserted into the ports for the two electrodes (Fig. S11). The chip was mounted on the microscope of a microfluidic station and the power electrode was connected to a high voltage amplifier, while the chip was grounded over the grounding electrode.

After an incubation of 12h, droplets containing cells, drug combinations and corresponding DNA-barcodes were transferred into a 3 ml syringe and injected through the droplet inlet port into a picoinjection chip (Fig. S11). Droplets were flushed at 180 µl h^-1^ and individual droplets were spaced out by injecting HFE oil with 1% PicoSurf surfactant at 700 µl h^-1^ to 1000 µl h^-1^ over the oil inlet. For cell lysis, barcode ligation and reverse transcription of mRNA released from lysed cells, we pico-injected a one-pot reaction mix containing 0.9% Igepal (Sigma Aldrich), 3x ligation buffer (NEB), 60,000 U/µl T4 Ligase (NEB), 1.5 mM dNTPs (ThermoFisher), 7.5 µM Template Switching Oligonucleotide (IDT), 12 U/µl Maxima -H reverse transcriptase (ThermoFisher) and 6,000 U/µl NxGen RNase Inhibitor (Lucigen). Flow rates for the reagent mix were adjusted according to the droplet frequency and size in order to inject the equivalent of ⅓ of the final droplet volume (Movie S3). In order to achieve the injection of reagents into the droplets passing by the injector nozzle, we applied a continuous electrical field of 0.1V using a function generator (Rigol). Pico-injection was performed over ∼1h during which all droplets (injection and collection) were kept on ice. Subsequently the emulsion was kept at RT for 30 min and then incubated for 90 min at 42 °C.

### Library preparation and sequencing

Upon reverse transcription of mRNA, all cDNA was barcoded according to drug treatments, and therefore, we broke the emulsion by adding 0.5 ml to 1 ml of 1H,1H,2H,2H-Perfluorooctanol (Abcr). The supernatant was transferred into a fresh Eppendorf tube, supplemented with 1x the volume of C1 dynabeads (ThermoFisher) at 2.5 µg µl^-1^ in 6x SSC buffer (ThermoFisher), and incubated at RT for 20 min. Beads were washed 2x in TE-SDS (10 mM Tris pH 8.0, 1 mM EDTA and 0.5% SDS) and 2x in nuclease free water (ThermoFisher). Beads were resuspended at 5 µg µl^-1^ followed by MseI (NEB) digestion according to the manufacturer’s instructions. The supernatant was purified 2x using SPRIselect beads (BD), first at 0.6x and then at 0.8x the volume of cDNA and finally amplified in KAPA HiFi ready mix (Roche) with 0.8 µM of SMART primer (Table S2) over a total of 13 cycles (PCR program in Table S3). Products were purified on 0.6x the volume SPRIselect beads and then analyzed using high sensitivity DNA chips on a 2100 Bioanalyzer (Agilent). Fragmentation of cDNA was performed to shorten the fragments and to introduce linker sequences. This was achieved using a Tn5-based tagmentation protocol for 3’ end libraries developed in house^35^. Fragments were amplified using a P5-SMART primer (Table S2) and an i7 indexed P7 adapter primers (Illumina, Table S2) at 0.75 µM in KAPA HiFi ready mix (PCR program table S4). Fragments were purified on 1x the volume of SPRIselect beads and size distributions were determined using a Bioanalyzer. All replicates were pooled at equimolar ratios and sequenced on a NextSeq 500 (Illumina) machine together with 10% PhiX spike-ins. Paired-end sequencing was performed by sequencing the barcode combination (Read 1, 26 bp) using the sequencing custom primer (Table S2) and the mRNA (Read 2, 59 bp).

### Statistics

In boxplots the center line represents the measured median and the upper box and lower box hinges corresponds to the first and third quartiles. The whiskers extending from the lower and upper hinges of the box represent the 1.5-fold interquartile range. The dots with the lines shown in the violin plot in Fig. 2d correspond to the mean with standard deviation for each of the measured cross contaminations.

### Gene expression data preprocessing

We used the SCANPY pipeline^36^ for gene expression preprocessing and quality control. Samples with low gene count and high ratio of mitochondrial genes (>15 percent) and genes with high dropout rate were filtered out. Read counts were normalized based on sequencing depth and z-score transformed. Batch effect (replicates) was removed by using the *combat* function of SCANPY. For dimension reduction we used Principal Component Analysis, followed by t-distributed stochastic neighbor embedding (TSNE)^37^.

### Clustering based quality control of gene expression data

To analyse the clustering of samples based on the different factors (Autosampler Drug, Braille Valves Drug, Combination) we used silhouette score analysis. Silhouette coefficient (b − a) / max(a,b) was calculated for each sample, where a was the mean intra- and b was the mean nearest-cluster distance. For each clustering factor, the average of Silhouette Coefficients were calculated (scikit-learn Python library). As silhouette score is dependent on the number of clusters, we created random clusters by permuting sample labels, thus cluster membership.

### Functional genomic analysis of gene expression signatures

Pathway activities were calculated using PROGENy method^17-19^. Individual drug specific pathway activities were calculated by fitting a linear model (pathway_activity ∼ YM155 + Imatinib + Trametinib).

To compare similarity between gene expression signatures from the high-throughput screen and LINCS-L1000 dataset^8^ we calculated consensus signatures for each drug of the high-throughput screen. To calculate consensus signatures, we fitted a linear model (gene_expression ∼ Drug_1_ + Drug_2_ + … + Drug_n_) for each gene of the expression matrix, and used the linear model coefficients as drug specific signature. To compare signature similarities, we calculated Spearman’s rank correlation coefficient between the drug signatures of high-throughput screen and LINCS-L1000 signatures. The similarity values were used as predicted values for ROC analysis, while true positives were the matched drug pairs between high-throughput screen and LINCS-L1000.

For cell viability predictions we used the CEVIChE method. CEVIChE predicts cell viability from gene expression changes based on a linear model, trained on a large compedion of matched cell viability and gene expression dataset^18^. As the measured genes of the high-throughput screen showed low overlap with the genes used by the original CEVIChE model, we retrained CEVIChE using only the genes measured in the high-throughput screen. This retrained CEVIChE model showed comparable performance (Pearson correlation between predicted and observed cell viability: 0.31) to the original method.

### Plate-based viability measurements to validate hits from microfluidic screen

Drug-plates were prepared in advance in a 4×4 checkerboard for each combination, such that after addition of cells, each drug was present at its GR35 concentration and a four-fold dilution series thereof. Each plate also contained media and DMSO negative controls and monotherapies for each drug. K562 cells were passaged the day before each experiment. On the day of the experiment, cells were washed once with PBS, then resuspended in FreeStyle 293 media (ThermoFisher), containing 1% FBS. Cells were added using the multistep function of a multichannel pipette to each pre-prepared drug-plate, such that each well had 200 µL final volume and approximately 2×10^4^ cells. The reservoir from which to aspirate cells was frequently refilled with freshly resuspended stock solution to ensure that cells remained in suspension. Plates were sealed with a gas-permeable foil (Sigma) and incubated for 48 hours. To prevent evaporation, plates were kept in the incubator within a box with ∼1 cm water, within the box the plates rested on tip boxes. After incubation, 22 µL PrestoBlue (ThermoFisher) cell viability reagent was added to each well and plates were resealed and returned to the incubator for 1 hour. Plates were then read using a Tecan microplate reader with excitation/emission wavelengths of 535/615 nm (20 and 10 nm wavelength bandwidth respectively). Based on the measured cell viability for monotherapies, we calculated expected cell viabilities using the Bliss independence model^21^ for each combination, for each concentration pair. The difference between expected and measured cell viability for combinations was averaged across all concentrations and was given as synergy score.

### Data and software availability

The datasets of the RNA-Sequencing experiments are available through GEO (accession GSE174696). Microfluidic control software and CAD designs of the chips can be downloaded from www.epfl.ch/labs/lbmm/downloads/mathur. The code used to analyze the transcriptomic data can be downloaded from https://github.com/bence-szalai/Combi-Seq-analysis.

