## Supplementary Information for "Combi-Seq: Multiplexed transcriptome-based profiling of drug combinations using deterministic barcoding in single-cell droplets"

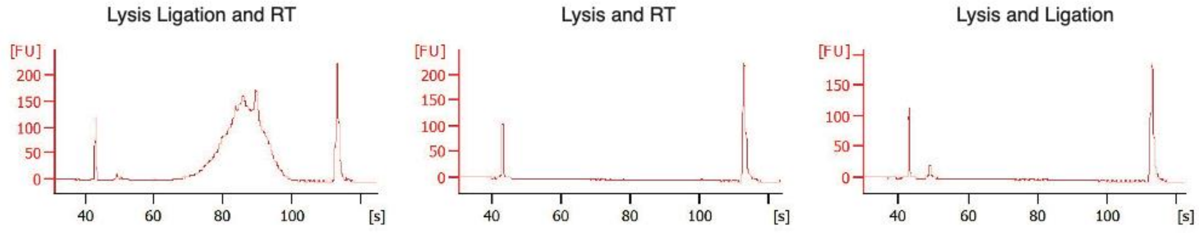


**Fig. S1:** Bioanalyzer traces showing specificity of ligation for the generation of functional barcodes. After picoinjections with the whole reaction mix (cell lysis, ligation and RT, left plot), reagents only for cell lysis and RT (center plot) and reagents for cell lysis and ligation (right plot), cDNA was purified and amplified (see methods) and loaded on a high sensitivity chip in a Bioanalyzer (Agilent).

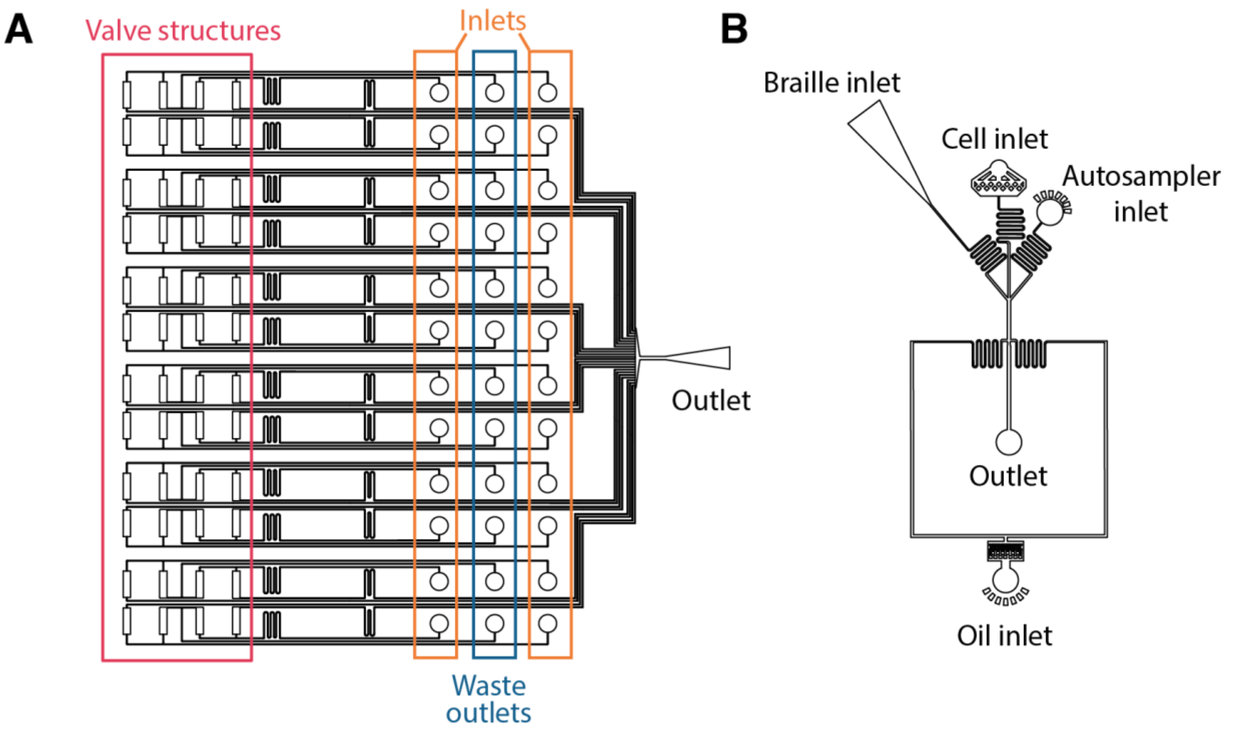


**Fig. S2: (A)** Chip design for the Braille valves. The chips were mounted on top of a Braille display with the rectangular valve structures on top of the pins. Inlets were used to inject drugs which in the default mode were directed to the waste outlets. Drug plugs were generated at the outlet to which a delay tube was connected horizontally into the funnel like structure. **(B)** Droplet maker chip design used to inject drug plugs via the horizontal Braille inlet and drugs from the 96-well plate via the Autosampler inlet. Cells were injected via the central cell inlet and co-encapsulated into droplets at the flow focusing junction by co-injecting oil in the perpendicular channels.

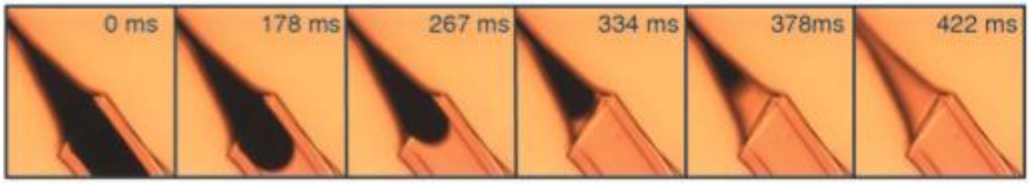


**Fig. S3:** Frames from a video sequence recording the injection of plugs labelled with Trypan Blue. The horizontal inlet port prevents plug breakups getting stuck at the inlet, which are then picked up by subsequent plugs causing cross-contaminations.

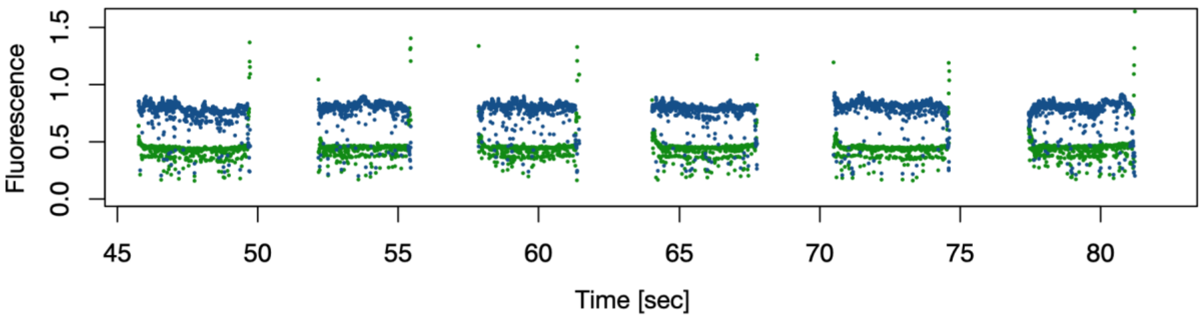


**Fig. S4**: Fluorescence intensities of six combinations generated with green (autosampler) and UV (Braille valves). UV fluorescence increased when plugs were injected and decreased at each end of the plug. Green signals decreased at the beginning of each plug since the continuously injected compounds from the autosampler were diluted. At the end of each plug the intensities increased, due to the end of the plug resulting in a higher concentration of the autosampler compound. Data was filtered for blue positive data points to remove green signals from between injected plugs.

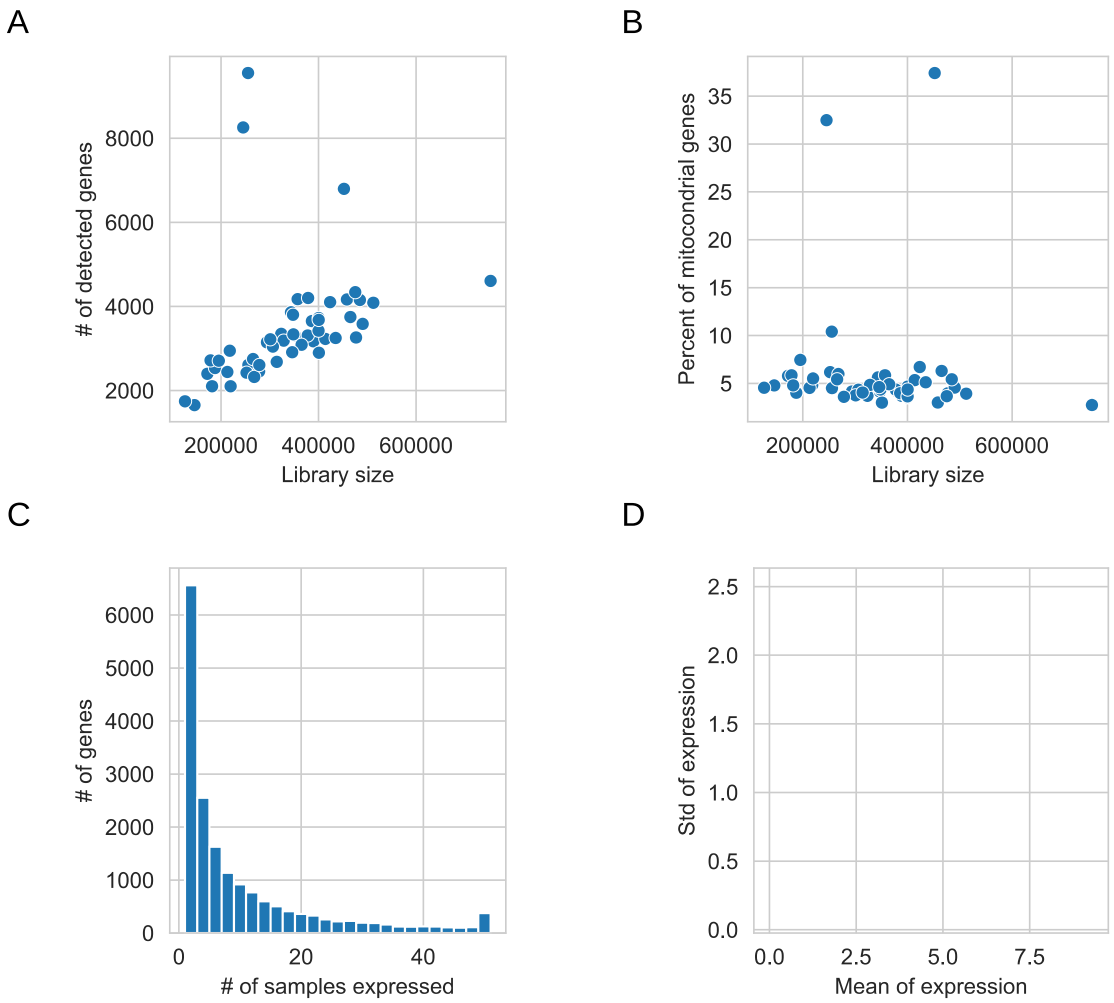


**Fig. S5**: Quality control for small scale drug screen **(a)** Relationship between the total counts / samples and the number of detected genes / samples. **(b)** Relationship between the total counts / sample and the percent of mitochondrial genes. **(c)** Distribution for the number of samples where a given gene was expressed. **(d)** Mean - Standard deviation relationship for the log1p transformed read counts.

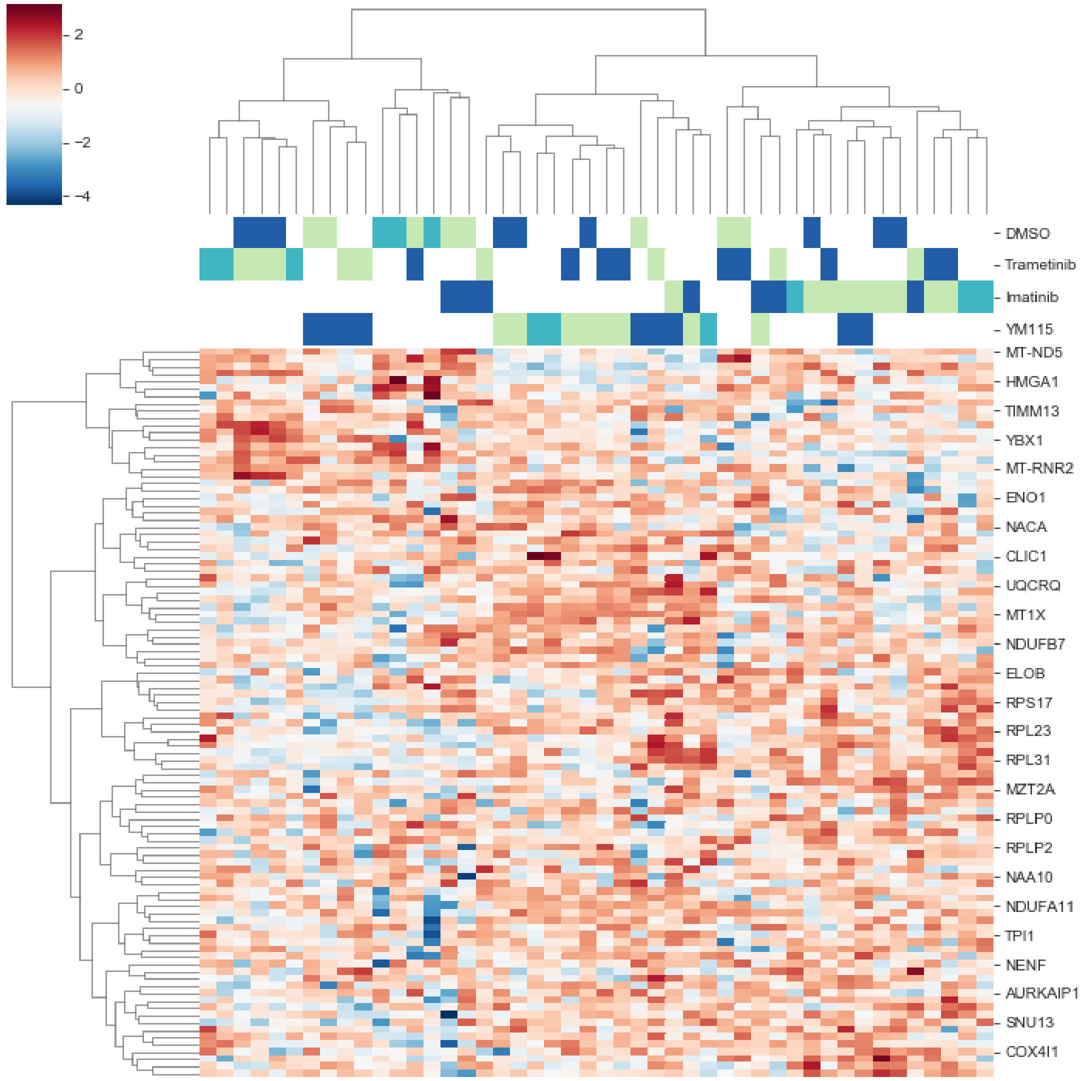


**Fig. S6**: Clustering of the small-screen samples based on the top 100 highly expressed genes. Normalized gene expression values (heatmap color code) were used to perform hierarchical clustering both on genes (x axis) and samples (y axis). Drugs of combinations are color coded (yellow: autosampler drug, blue: braille valves drug, cyan both drugs).

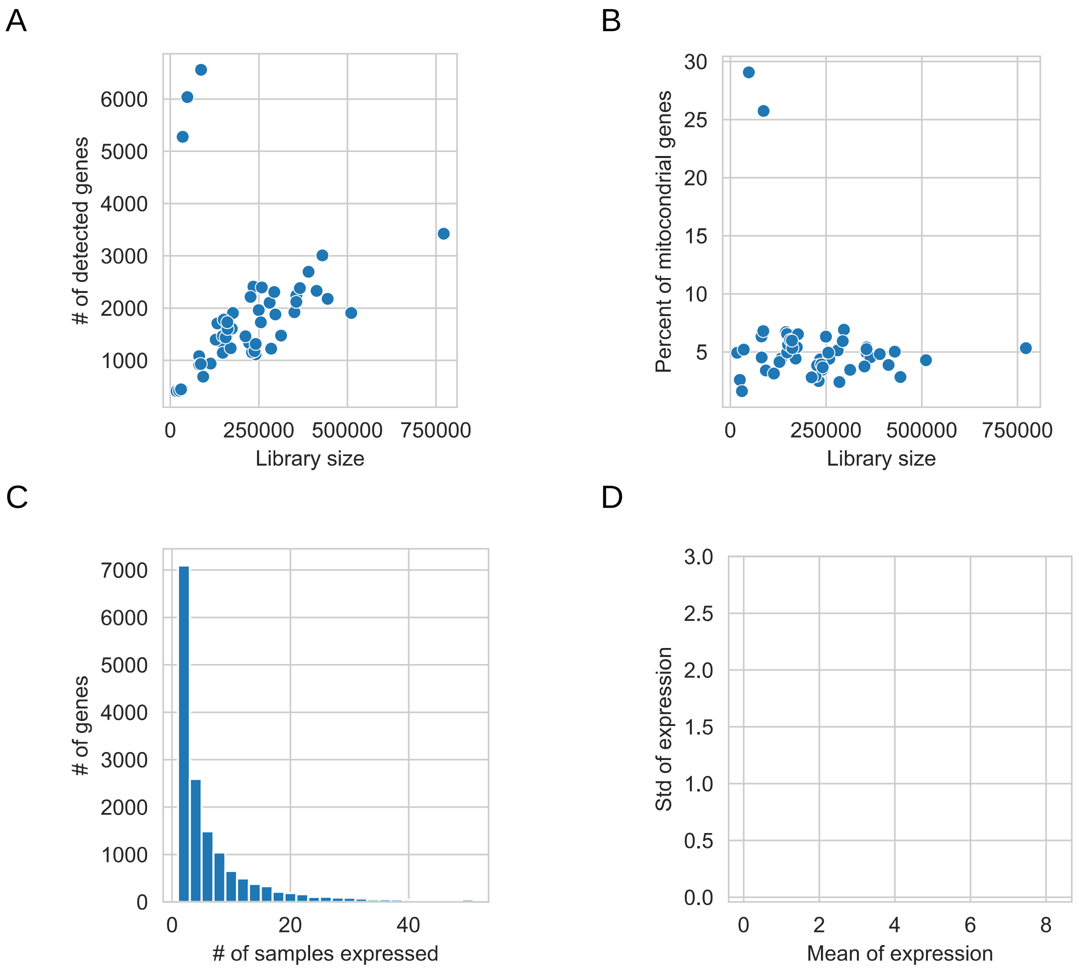


**Fig. S7**: Quality control for small scale drug screen with swapped barcoding mode: **(a)** Relationship between the total counts / samples and the number of detected genes / samples. **(b)** Relationship between the total counts / sample and the percent of mitochondrial genes. **(c)** Distribution for the number of samples where a given gene was expressed. **(d)** Mean - Standard deviation relationship for the log1p transformed read counts.

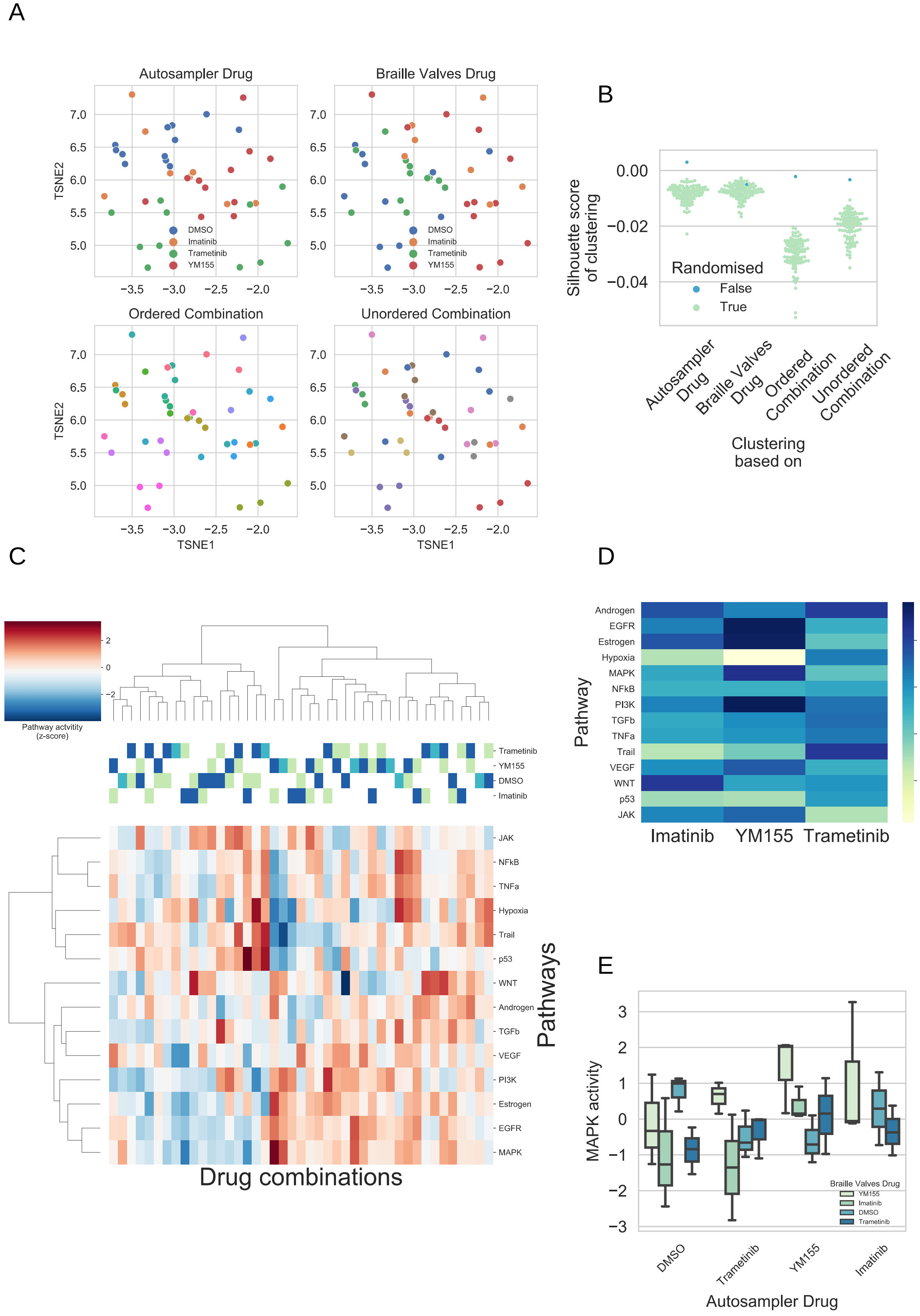
A
Autosampler Drug
Braille Valves Drug
Ordered Combination
Unordered Combination
TSNE1
TSNE2
DMSO
Imatinib
Trametinib
YM155
B
Silhouette score of clustering
Randomised
False
True
Autosampler Drug
Braille Valves Drug
Ordered Combination
Unordered Combination
Clustering based on
C
Pathway actvitity (z-score)
Trametinib
YM155
DMSO
Imatinib
JAK
NFkB
TNFa
Hypoxia
Trail
p53
WNT
Androgen
TGFb
VEGF
PI3K
Estrogen
EGFR
MAPK
Drug combinations
Pathways
D
Pathway
Androgen
EGFR
Estrogen
Hypoxia
MAPK
NFkB
PI3K
TGFb
TNFa
Trail
VEGF
WNT
p53
JAK
Imatinib
YM155
Trametinib
Model coefficient
E
MAPK activity
Braille Valves Drug
YM155
Imatinib
DMSO
Trametinib
DMSO
Trametinib
YM155
Imatinib
Autosampler Drug

**Fig. S8** Functional analysis for small scale drug screen with swapped barcoding mode **(a)** TSNE plots of normalized gene expression data. Samples are color coded based on Autosampler Drug (top left panel), Braille Valves Drug (top right panel), Ordered Combination (bottom left panel) and Unordered Combination (bottom right panel). Color code is labeled for Autosampler and Braille Valves Drugs (top panels). **(b)** Silhouette scores of sample clustering based on Autosampler / Braille Valves Drugs and Ordered / Unordered Combinations. Silhouettes scores are compared to random distributions (color code) created by permuting sample labels. **(c)** Pathway activity heatmap of samples. PROGENy pathway activities were calculated for each sample (z-scores of pathway activities, color code) and the pathway activity matrix was hierarchically clustered. Drugs of combinations are color coded (yellow: autosampler drug, blue: braille valves drug, cyan both drugs). **(d)** Drug induced pathway activity changes. Linear model (pathway_activity ~YM155 + Imatinib + Trametinib) was fitted for each pathway, and the linear model coefficients (color code) for each drug is plotted as a heatmap. (e) Drug induced MAPK activity changes. MAPK activity (y axis) grouped based on Autosampler Drug (x axis) and Braille Valves Drug (color code) and plotted as a boxplot.

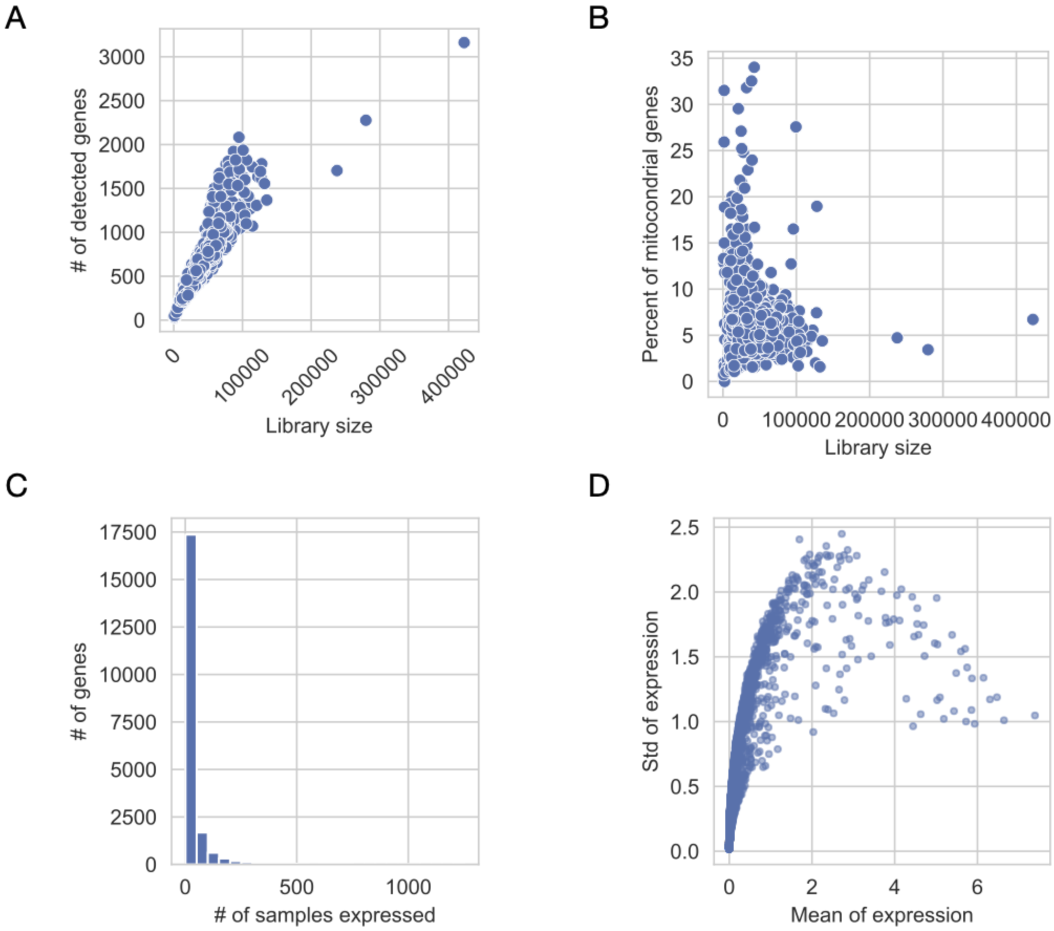


**Fig. S9:** Quality control for large scale drug screen **(a)** Relationship between the total counts / samples and the number of detected genes / samples. **(b)** Relationship between the total counts / sample and the percent of mitochondrial genes. **(c)** Distribution for the number of samples where a given gene was expressed. **(d)** Mean - Standard deviation relationship for the log1p transformed read counts.

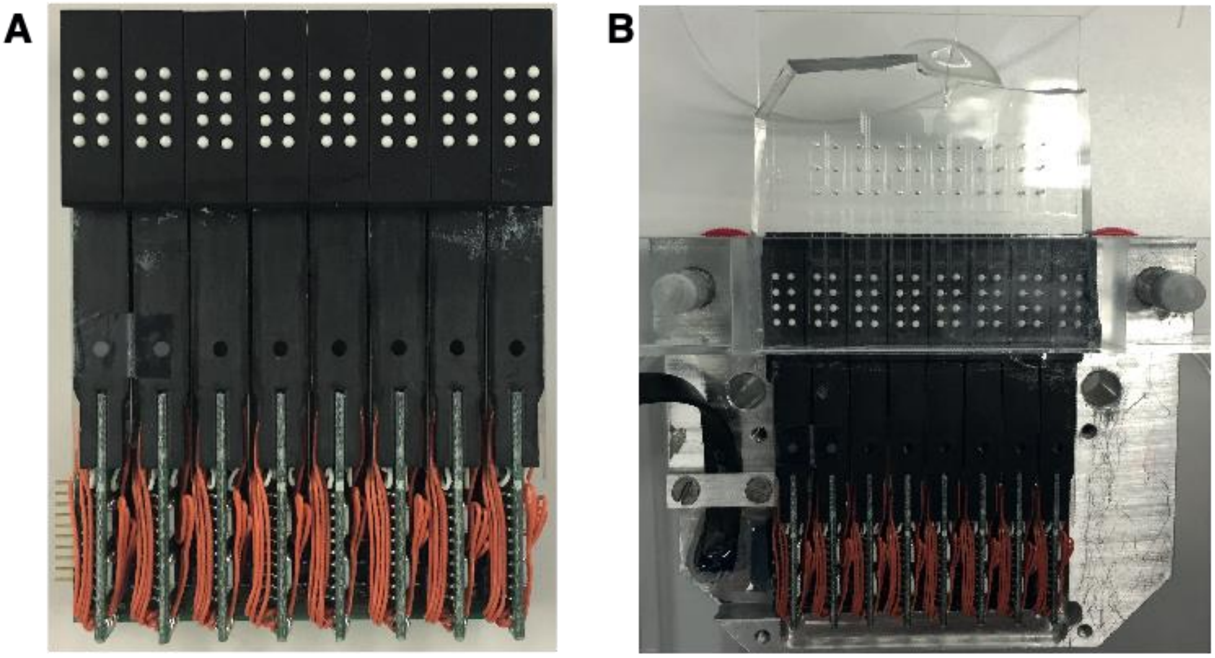

**Fig. S10**: Pictures of the Braille display. (A) Braille display alone. (B) Braille display mounted on the home-made chip holder. A Braille valve chip was mounted on top so that the Braille pins were aligned with the rectangular valve structures of the chip.

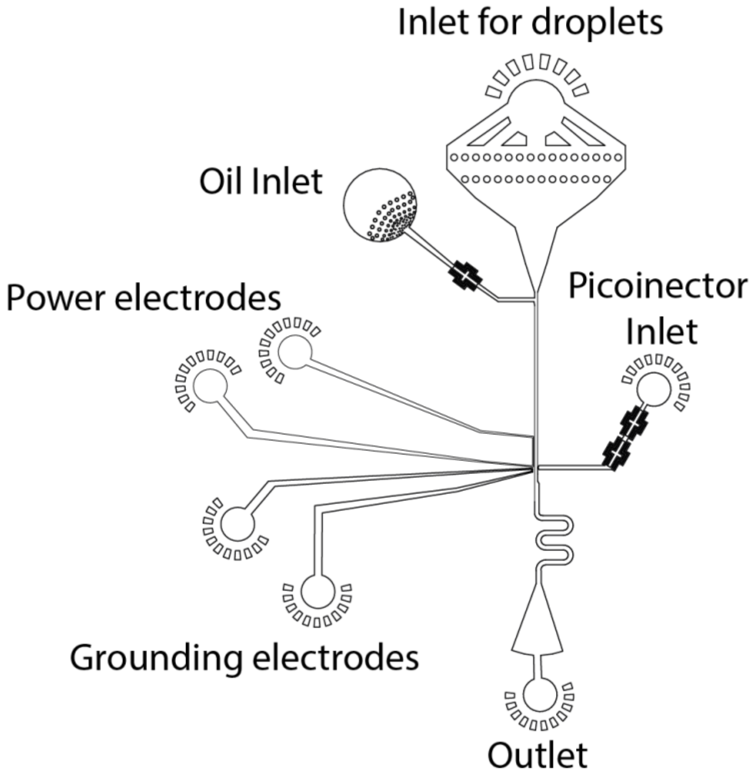


**Fig S11:** Chip design used to manufacture molds for Pico-injection devices. Channels for the power and grounding electrodes were filled with solder and cables to connect the chip with a function generator or for grounding were inserted into the inlet ports.

Table S1

| | Chip 1 | Chip 2 | Chip 3 |
|---|---|---|---|
| Average contamination UV channel | 1.44% (n=99) | 0.98% (n=80) | 2.3% (n=171) |
| Average contamination green channel | 0.92% (n=99) | 0.47% (n=80) | 0.73% (n=171) |

Table S2

| Sequence Name | Sequence (5’ -> 3’) |
|---|---|
| TSO | AAGCAGTGGTATCAACGCAGAGTGAATrGrGrG |
| SMART-Primer | AAGCAGTGGTATCAACGCAGAGT |
| Tn5ME (loaded on Tn5) | GTCTCGTGGGCTCGGAGATGTGTATAAGAGACAG |
| Tn5MErev | [phos]CTGTCTCTTATACACATCT |
| i7 indexed P7 adapter primers | CAAGCAGAAGACGGCATACGAGATnnnnnnnnGTCTCGTGGGCTCGG |
| P5-SMART adapter primer | AATGATACGGCGACCACCGAGATCTACACGCCTGTCCGCGGAAGCAGTGGTATCAACGCAGAGTAC |
| Custom Sequencing Primer | GCCTGTCCGCGGAAGCAGTGGTATCAACGCAG AGTAC |

Table S3

| Whole transcriptome amplification | | |
|---|---|---|
| **Step** | **Temperature** | **Time** |
| Initial denaturation | 95 °C | 3 min |
| 4 cycles | 98 °C<br>65 °C<br>72 °C | 20 sec<br>45 sec<br>3 min |
| 9 cycles | 98 °C<br>67 °C<br>72 °C | 20 sec<br>20 sec<br>3 min |
| Final extension | 72 | 5 min |
| Hold | 4 °C | |
| **Tagmentation PCR** | | |
| **Step** | **Temperature** | **Time** |
| Initial denaturation | 95 °C | 30 sec |
| 12 cycles | 98 °C<br>58 °C<br>72 °C | 20 sec<br>15 sec<br>30 sec |
| Final extension | 72 | 3 min |
| Hold | 10 °C | |

Table S4

| Drug | Conc. [µM] | Drug Class | Targets | ChEMBL ID | Provider |
|---|---|---|---|---|---|
| 10Z-Hymenialdisine | 4.6088 | Kinase-Inhibitor | MEK1 | CHEMBL361708 | TOCRIS |
| 5-Iodotubercidine | 6.6510 | Anti-metabolite | ADK/INSR/ PKA/CK1 | CHEMBL99203 | Selleckchem |
| AT9283 | 0.1718 | Kinase-Inhibitor | AURKA/AURKB/JAK2/JAK3 | CHEMBL495727 | Selleckchem |
| Baricitinib | 1.5004 | Kinase-Inhibitor | JAK1/JAK2 | CHEMBL2105759 | Selleckchem |
| Blebbistatin | 8.9951 | Cytoskelleton | MYH2 | CHEMBL1328324 | Selleckchem |
| Clofarabine | 0.8862 | Anti-metabolite | RRM1 / DNA Polymerase | CHEMBL1750 | Selleckchem |
| Cytarabine | 100 | Anti-metabolite | DNA / RNA Polymerase | CHEMBL803 | Selleckchem |
| Dacarbazine | 3.5754 | Alkylating Agent | DNA Antimetabolite | CHEMBL476 | Selleckchem |
| Decitabine | 1.6615 | Anti-metabolite | DNMT1 | CHEMBL1201129 | Selleckchem |
| Dovitinib | 0.1592 | Kinase-Inhibitor | FLT3/c-Kit/FGFR1/FGFR3 | CHEMBL522892 | Selleckchem |
| Doxorubicin | 0.5333 | Anthracycline | TOP2 | CHEMBL53463 | Selleckchem |
| Epirubicin | 0.1166 | Anthracycline | TOP2 | CHEMBL1200981 | Selleckchem |
| Fludarabine Phosphate | 100 | Anti-metabolite | DNA Antimetabolite | CHEMBL1096882 | Selleckchem |
| Fluoruracil | 7.8542 | Anti-metabolite | TYMS | CHEMBL185 | Selleckchem |
| Gemcitabine | 0.0437 | Anti-metabolite | DNA Antimetabolite | CHEMBL888 | Selleckchem |
| Gimeracil | 100 | | DPYD | CHEMBL1730601 | Selleckchem |
| H-7 dihydrochloride | 75.6496 | Kinase-Inhibitor | PRKC/PKG/PKA | | TOCRIS |
| Hematoxylin | 2.5893 | | EGFR/ERBB2/c-MET/c-KIT/SRC | CHEMBL477197 | Selleckchem |
| Imatininb | 0.2368 | Kinase-Inhibitor | BCR-ABL | CHEMBL941 | Selleckchem |
| Methotrexat | 100 | Kinase-Inhibitor | DHFR | CHEMBL34259 | Selleckchem |
| Mitomycin C | 0.9501 | DNA-crosslinker | DNA Synthesis | CHEMBL105 | Selleckchem |
| NMS1286937 (Onvansertib) | 0.1228 | Kinase-Inhibitor | PLK1 | CHEMBL1094408 | Selleckchem |
| Olomoucine | 4.9738 | Kinase-Inhibitor | CDK2/MAPK3 | CHEMBL280074 | TOCRIS |

| | | | | | |
|---|---|---|---|---|---|
| Oxaliplatin | 10.1424 | Alkylating Agent | DNA | CHEMBL414804 | Selleckchem |
| PF-562271 | 8.2299 | | FAK | CHEMBL1084546 | Selleckchem |
| Razoxane | 13.8055 | Anthracycline | TOP2 | CHEMBL1738 | Selleckchem |
| SB747651A | 0.8732 | Kinase-Inhibitor | MSK1/MSK2 | CHEMBL188434 | TOCRIS |
| SF-1126 | 3.2887 | Kinase-Inhibitor | PI3K/mTOR | CHEMBL2326966 | Santa Cruz |
| Sonolisib | 4.7513 | Kinase-Inhibitor | PI3K | CHEMBL411907 | Abcam |
| Streptozotocin | 100 | | DNA | CHEMBL1603 | Selleckchem |
| Sunitinib Malate | 7.8137 | Kinase-Inhibitor | VEGFR2/PDGFRb | CHEMBL1567 | Selleckchem |
| Tabloid (Thioguanine) | 0.9472 | Anti-metabolite | DNMT1 | CHEMBL727 | Selleckchem |
| Trametinib | 0.2085 | Kinase-Inhibitor | MEK1/2 | CHEMBL2103875 | Selleckchem |
| Triciribine | 8.3030 | Kinase-Inhibitor | AKT1/AKT2/AKT3 | CHEMBL462018 | Selleckchem |
| Wortmannin | 1.5465 | Kinase-Inhibitor | PI3K | CHEMBL428496 | Selleckchem |
| YM155 | 0.0003 | | BIRC5 | CHEMBL2110734 | Selleckchem |
